# Navigating Memorability Landscapes: Hyperbolic Geometry Reveals Hierarchical Structures in Object Concept Memory

**DOI:** 10.1101/2024.09.22.614329

**Authors:** Fiona M. Lee, Marc G. Berman, Andrew J. Stier, Wilma A. Bainbridge

## Abstract

Why are some object concepts (e.g., birds, cars, vegetables, etc.) more memorable than others? Prior studies have suggested that features (e.g., color, animacy, etc.) and typicality (e.g., robin vs. penguin) of object images influences the likelihood of being remembered. However, a complete understanding of object memorability remains elusive. In this study, we examine whether the geometric relationship between object concepts explains differences in their memorability. Specifically, we hypothesize that image concepts will be geometrically arranged in hierarchical structures and that memorability will be explained by a concept’s depth in these hierarchical trees. To test this hypothesis, we construct a Hyperbolic representation space of object concepts (N=1,854) from the THINGS database (Hebart et al., 2019), which consists of naturalistic images of concrete objects, and a space of 49 feature dimensions derived from data-driven models. Using ALBATROSS (Stier, A. J., Giusti, C., & Berman, M. G., In prep), a stochastic topological data analysis technique that detects underlying structures of data, we demonstrate that Hyperbolic geometry efficiently captures the hierarchical organization of object concepts above and beyond a traditional Euclidean geometry and that hierarchical organization is related to memorability. We find that concepts closer to the center of the representational space are more prototypical and also more memorable. Importantly, Hyperbolic distances are more predictive of memorability and prototypicality than Euclidean distances, suggesting that concept memorability and typicality are organized hierarchically. Taken together, our work presents a novel hierarchical representational structure of object concepts that explains memorability and typicality.

## Introduction

Among the vast array of images that we encounter each day, how do our brains decide what to remember and what to forget? While there is a large degree of individual variation in what is remembered [1,2], there is also surprisingly high consistency across people in their memories: certain stimuli are intrinsically remembered better than others [3,4]. Memorability, a stimulus property that measures how likely an item is going to be remembered across people, can predict people’s memory with high success across tasks [5,6,7], participants [8], and stimulus categories [3,4,9]. However, there has been limited success in understanding what makes something memorable, even when examining an exhaustive set of object images and their features [10]. Here, we propose that we have been limited by traditional assumptions of memory being organized in a Euclidean space, with restricted interpretability. However, by examining an entirely different cognitive geometry—a Hyperbolic space— we can more directly form intuitions about what makes something memorable and about the organization of our cognitive spaces more broadly.

Prior approaches aiming to understand memorability have been unable to explain why certain stimuli are more memorable than others. Several studies have shown that memorability is not a proxy for some singular trait, such as aesthetic value, emotion, visual saliency, color, or ratings of typicality [4,11]. Other work has tested combinations of multiple visual and semantic features (e.g., animacy, redness, naturalness, etc.) to capture higher levels of variance in memorability (e.g., 61.66% of the noise ceiling using 49 features in [10]), but these approaches have failed to reveal relationships between features and memorability that can generalize beyond those images and features.

Still other work has modeled images as points within a multidimensional representation space, with axes representing stimulus features, to examine how memorability relates to an item’s location within that feature space [12,13,14]. Decades of prior work have suggested that memorability may be closely tied to the typicality of an item [15,16,17,18], which can be represented as whether that item has many other items that are close by (and therefore similar to it) in a multidimensional feature space. For example, a *horse* shares similarity with more animals than a *sea urchin* does, and thus the former usually is considered as more prototypical than the latter. However, there has been mixed evidence on whether memorable items are more atypical [13,17] or prototypical [15,16,18], with recent evidence showing mild evidence in favor of more prototypical images being more memorable [10]. This may be due to the fact that most prior works focused only on the within-category aspect of typicality, namely, how typical an item is compared to all other items in the same category. Here, we introduce another definition of typicality to capture between-category relationships. Specifically, we measure how dissimilar an item is from all items outside of its category. Items that are more different from other categories are less likely to be falsely classified into other categories. We argue that in addition to the similarity within each category, the distinctiveness across categories also plays a role in human memory.

Rather than looking solely at prototypicality, another possibility is that memorability is related to an item’s location within a multidimensional feature space. Some work has investigated this, but from a Euclidean geometry perspective. Most studies examine image feature spaces as operating within a Euclidean geometric space. For example, one common approach is to find Euclidean embeddings that best capture pairwise similarity relationships between stimuli through multidimensional scaling (MDS) on human judgments. In turn, these embeddings are used to discover how features and the clustering of objects in the embedding space are related to memorability [10,12,13,17,18,19,20]. Other cognitive processes, such as the categorization of artificial and natural objects [21,22,23] and face recognition [12,24] have been largely studied in Euclidean feature spaces.

However, recent work from other fields has discovered that many biological stimuli and human judgments of them are better modeled within a Hyperbolic space, i.e., a space that acts as a continuous approximation to hierarchical tree structures. For example, plants and animals produce various odors as mixtures of different single molecule odors through hierarchical biochemical processes [25]. Similarly, patterns of gene expression [26] and representations of human and macaque visual perception [27,28] show a hierarchical nature that is better explained by a Hyperbolic geometry. One of the key differences between a Hyperbolic and Euclidean space is how distances are captured (Fig. 1). In a standard cube-like Euclidean space with no curvature, distances follow straight lines between any two points. In contrast, a shell-like Hyperbolic space simulates a tree structure, where the distance between two nodes approximates the path from both nodes back to their lowest common predecessor with a smooth curve. Therefore, as trees get deeper with more nodes, the distance between leaf nodes (i.e., those farther away from the origin) increases. This gives rise to another key characteristic of Hyperbolic spaces: they can generally preserve complex hierarchical structures with fewer dimensions than Euclidean spaces [29]. As a result, in a representational space where the distance between stimuli represents their similarity, Hyperbolic representations can expand the size of the tree-like structure to make room for more dissimilar stimuli within existing dimensions. In contrast, the only way to make room for increasingly dissimilar stimuli in a Euclidean space is to add more dimensions.

**Fig. 1:**
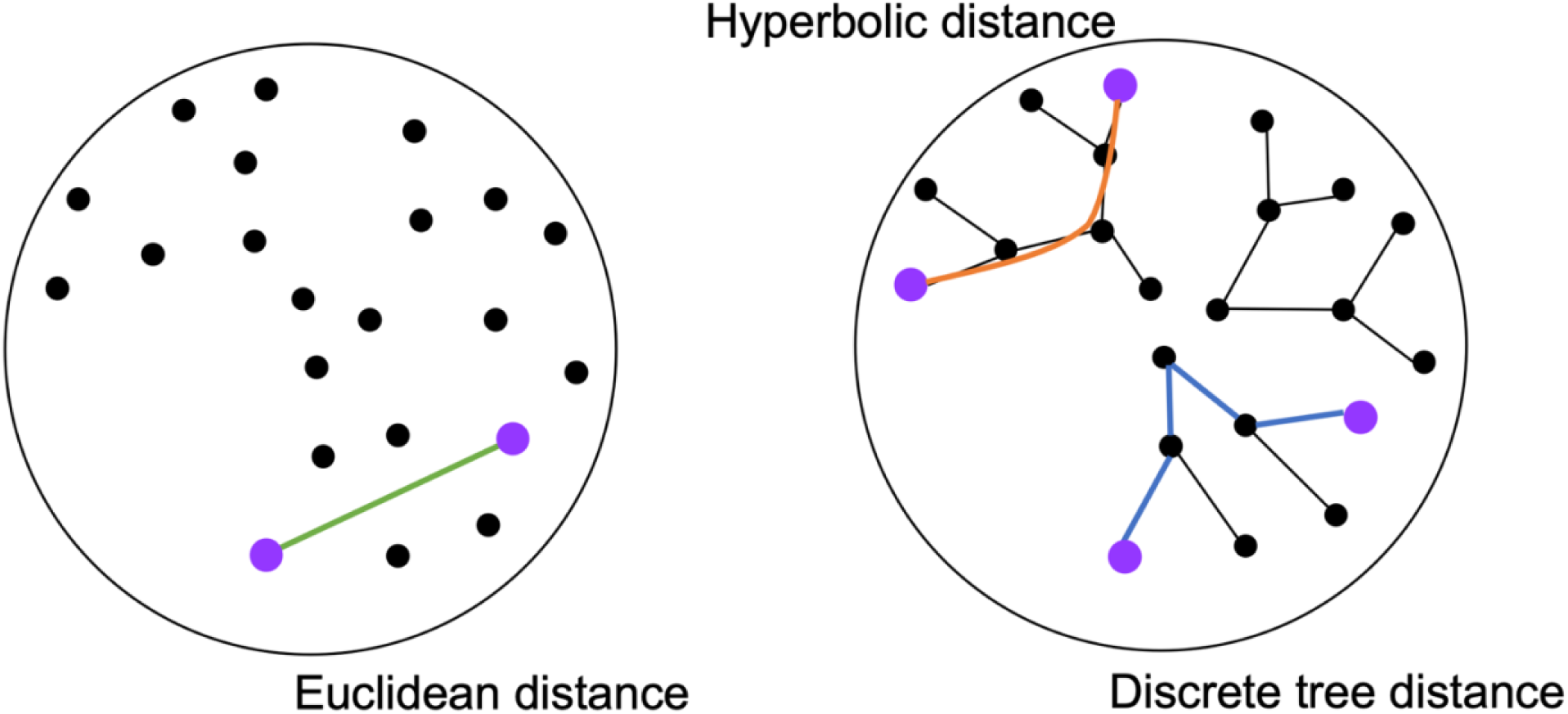
Euclidean distance (left), discrete tree distance (bottom-right), and Hyperbolic distance (upper-right) represented by the colored lines between two purple nodes. The Euclidean distance (green) between the same two nodes follows the shortest straight path. Hyperbolic geometry simulates a hierarchical structure, which can be conceptualized as trees with roots situated closer to the center and leaves growing outwards. In such a tree-like structure, the discrete tree distance (blue) between two points is measured by tracing back to their lowest shared root. Within a continuous space, the Hyperbolic distance (orange) approximates this discrete tree distance with a smooth curve, expanding the distance between two nodes relative to the Euclidean distance between the nodes.

We propose that cognitive feature spaces are also better captured by hierarchical tree-like networks. Prior work has shown that memories are integrated into network structures [30,31,32,33], in which related items are connected in memory, and we traverse that network to encode and retrieve items. For example, autobiographical and event memories are considered to be organized in hierarchical network structures, in which specific events are retrieved along top-down pathways [34,35,36,37]. In addition, within language-based semantic models, memorable words tend to be at the roots of a semantic network, while forgettable words tend to be at the leaves [8]. Until now, a similar pattern has not yet been tested with images, potentially because it has been unclear how to form a network structure capturing the many continuous features along which images vary. With the inherent hierarchical structure in Hyperbolic geometry, we can transform the image space into a continuous tree-like network. Such a structure allows us to test whether memorability may represent the position of an item within the whole tree, where more memorable items are hypothesized to be closer to the roots.

Here, we examined different geometric feature spaces of concrete object representations and the consequences of these spaces for understanding memorability. Using all 1,854 concrete objects from THINGS [38], a comprehensive object image database, we tested whether a Hyperbolic geometric space can better capture the relationship between the spatial organization of object representations and memorability than a Euclidean space does. In Hyperbolic space, we anticipate that memorable objects will cluster near the center of the space (roots) while forgettable objects scatter farther away from the center (leaves). Moreover, we expect to find a positive correlation between typicality and memorability in the Hyperbolic space. Most importantly, we expect that more variance in memorability could be explained in the Hyperbolic space than in the Euclidean space.

## Results

Our analysis (Fig. 2) starts by modeling the space of concrete object concepts in different geometries, through which we obtain the best-fitting Hyperbolic and Euclidean space models for our data. In these spaces, we locate object concepts and their features and examine how the organization of concepts differs in these two geometries. To answer our questions about the relationship between spatial location and memorability of concrete object concepts, we investigate the role of concept locations in explaining memorability in Hyperbolic and Euclidean spaces. Next, we examine how concept locations relate to within-category and between-category aspects of typicality. Finally, we address the question of the relationship between typicality and memorability.

**Fig. 2:**
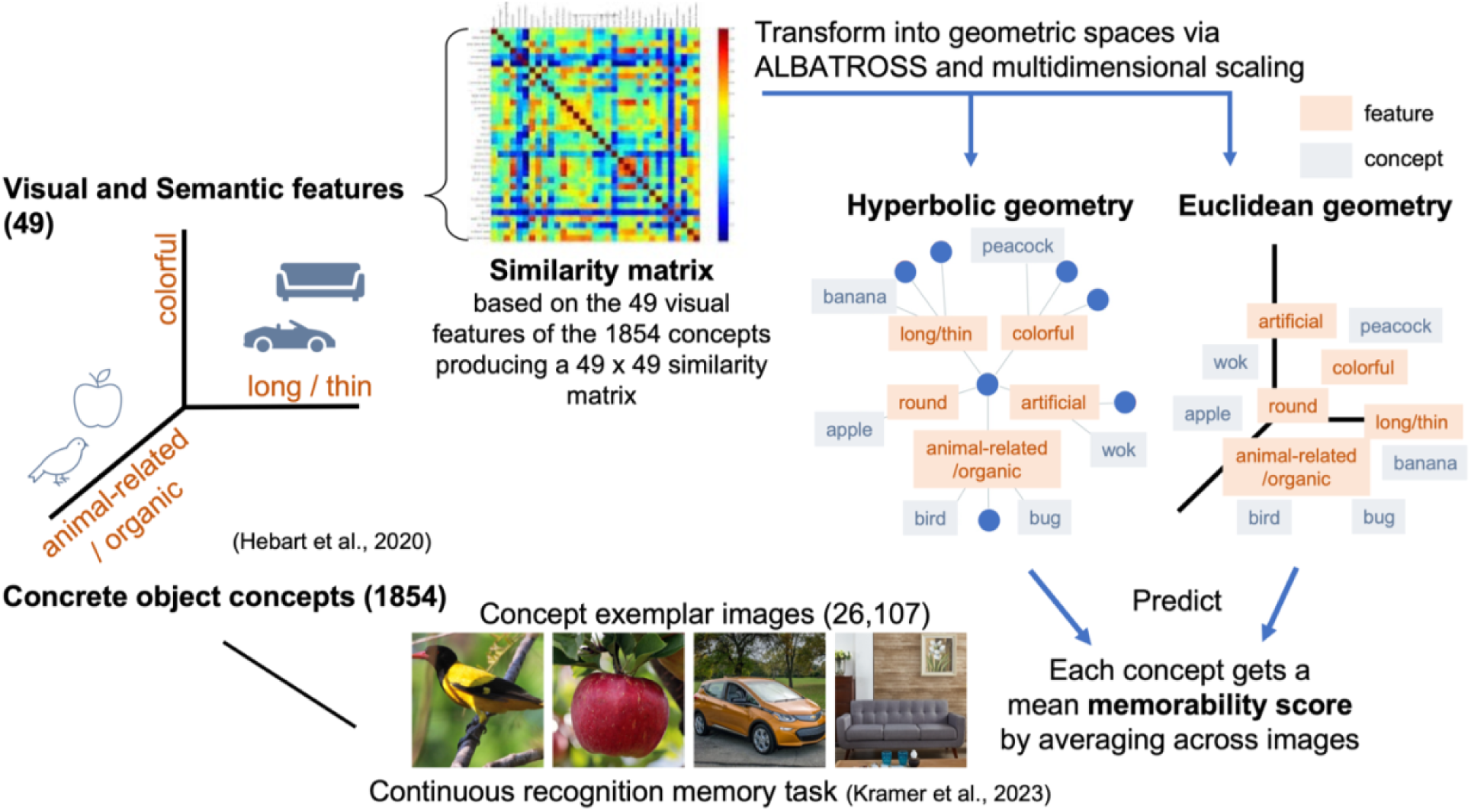
Overview of our analysis. Using all 1854 concepts and 49 visual and semantic features in the THINGS database (Hebart et al., 2020), we generated a similarity matrix and modeled the object spaces in Hyperbolic and Euclidean geometries, respectively (see more details in Methods). The spatial arrangements of concepts in these two spaces were used to predict the memorability scores of each concept, obtained in a continuous recognition task (Kramer et al., 2023).

### Hyperbolic geometry fits object image space with fewer dimensions

We first determined whether a Euclidean (flat) or Hyperbolic (curved) geometry can best describe the organization of object concept space. We expected to see that a Hyperbolic space model can fit our data and require fewer dimensions than a Euclidean space model does [29]. To test this hypothesis, we utilized the THINGS database, which contains images of most nameable concrete objects in the American English language [38,39]. The database contains photographs of 1,854 object concepts at a basic level of categorization (e.g., cat, apple). These object concepts are sorted into 27 broad high-level categories (e.g., animals, food) through data-driven and human-labeling approaches. Each object concept consists of on average 12 exemplar photographs (e.g., 12 cat photographs), resulting in 26,107 images in the database overall. Finally, each object concept is characterized by 49 features, which capture the perceptual (e.g., is white) and semantic properties (e.g., is animal-related) of concepts and were derived from odd-one-out triplet similarity ratings amongst the 1,854 object concepts. Thus, these features represent the latent dimensions people used to judge the similarity of concepts but do not directly capture visual features such as brightness or hue.

Via ALBATROSS (Stier, A. J., Giusti, C., & Berman, M. G., In prep), a statistical topological data analysis protocol, we determined the Hyperbolic and Euclidean spaces that best fit the 1,854 concepts. This protocol samples points from different geometries and compares resulting distance matrices to the empirical dissimilarity matrix. Specifically, we constructed an empirical dissimilarity matrix from correlations between features across object concepts; these correlations represent the degree to which features co-occur in pairs of objects. Then we sampled points from Euclidean unit cubes of different dimensions and 3-dimensional (3D) Hyperbolic shells with various maximum and minimum radii and compared the ensuing distance matrices to the empirical dissimilarity matrix. By performing a grid search we found values for the number of dimensions (for Euclidean cubes) and the minimum and maximum radii (for Hyperbolic shells) that minimized the difference between the distances sampled from these spaces and the empirical dissimilarity matrix.

We found that distance matrices from a 3-dimensional (3D) Hyperbolic shell and a 16-dimensional (16D) Euclidean unit cube were not significantly different from the empirical dissimilarity matrix in a chi-square test (*p_Hyperbolic_* = 1.0, *p_Euclidean_* = 0.405). This suggests that both geometries capture the spatial organization of similarity relationships among the 49 object features. However, the Hyperbolic space has a lower chi-square statistic (*X*^2^ (1, *N* = 2,401) ≈ 0.0) than the Euclidean space (*X*^2^ (1, *N* = 2,401) = 12.514), suggesting that the 3-dimensional Hyperbolic geometry better captures the empirical organization of features in concept space. In addition, we found that Hyperbolic models beyond 5 dimensions were statistically distinct from the empirical dissimilarity matrix (Supplementary Table 1) indicating that higher dimensional Hyperbolic geometries are not accurate when describing the feature space of the THINGS data. In contrast, a minimum of 16 Euclidean dimensions are required to capture the similarity relationships between concept features. This is in line with our prediction [29] that fewer Hyperbolic dimensions will be required to preserve the empirical dissimilarity relationships among the 49 features.

### Hyperbolic geometries better capture semantic relationships between and within categories

Having determined the overall geometry of features in concept space, we next examined how the features and concepts are organized differently in these spaces. We started by obtaining the locations of features and concepts in the best fit Hyperbolic and Euclidean spaces. Then, we made a number of observations and comparisons of the spatial organizations of features and concepts between these two spaces. Finally, we analyzed the difference in organization between spaces at the broader category level.

In order to determine the location of each feature, we mapped all feature embeddings from the 49-dimensional dissimilarity matrix to the best fit lower dimensional spaces through a regular Euclidean and a Hyperbolic multidimensional scaling approach (MDS, see Fig. 3). Within the Hyperbolic space, we expected the concepts to be organized hierarchically, in which concepts that belong to more general categories, such as “animals” and “food”, will be closer to the origin (i.e., the center of the space) and the roots of tree-like hierarchies; whereas concepts that are in more specialized categories, such as “clothing accessory”, would be further away from the origin as the leaves of hierarchical trees.

**Fig. 3:**
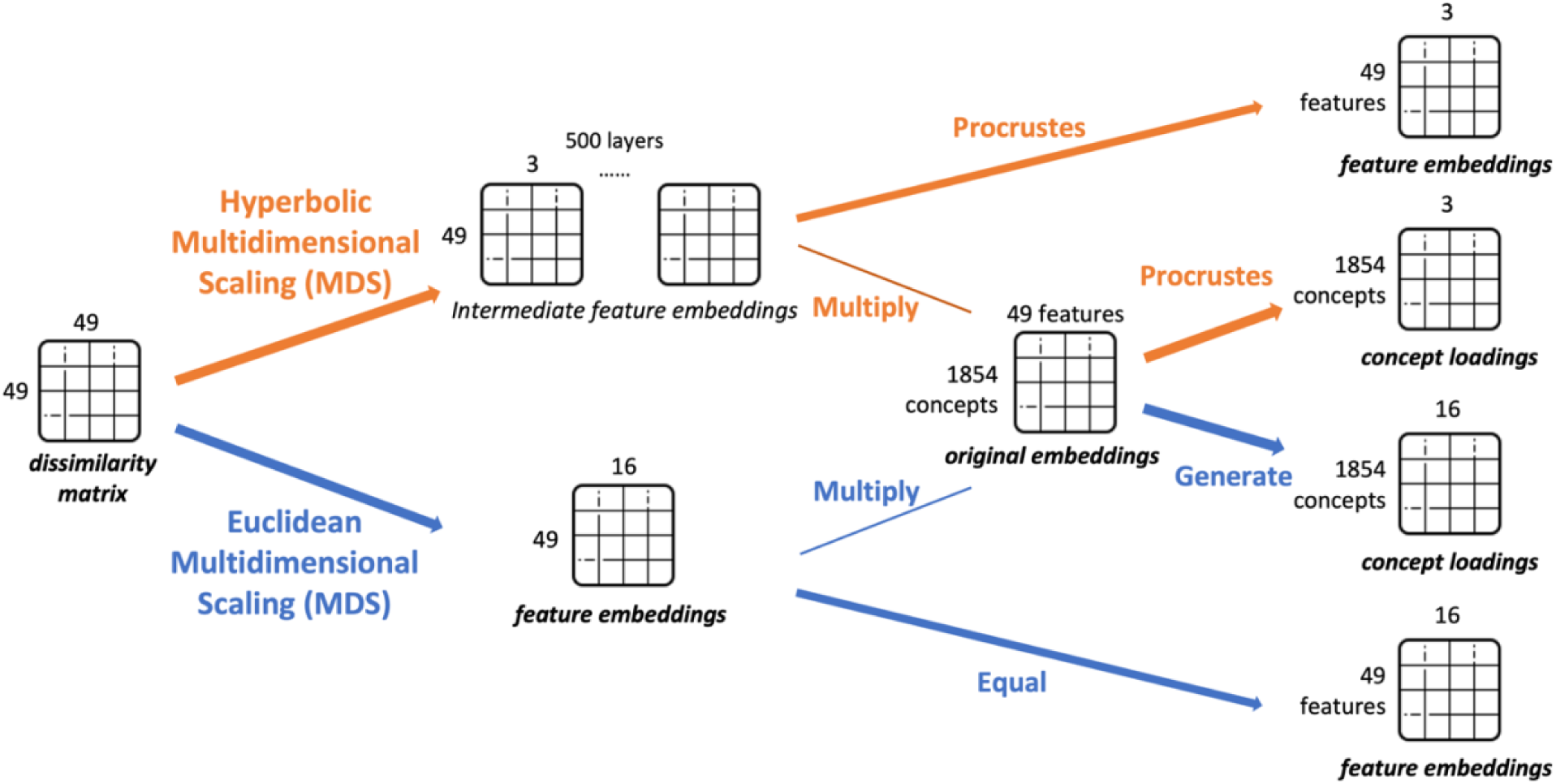
Placing 49 features and 1,854 concepts into 3D Hyperbolic and 16D Euclidean spaces. A dissimilarity matrix is created from the correlational distances between 49 features, from which the new embeddings of 49 features in the 16D Euclidean and the 3D Hyperbolic spaces are derived through regular Euclidean MDS or Hyperbolic MDS. The new embeddings of the 1,854 concepts are obtained by multiplying the original embeddings with the new feature embeddings. Note that the Hyperbolic MDS generates 500 layers of intermediate feature embeddings through a stochastic process, so the new embeddings of features and concepts in the 3D Hyperbolic space are finalized via Procrustes transformation. See more details in Methods.

Examining the embeddings in 3D Hyperbolic space, we observe that the 49 features are situated at various radii (distance from the origin of the space) around the surface of the inner empty core, defined by the space’s minimum radius of 8.4 arbitrary units (a.u.) (Fig. 4). The features have an average radius of 8.71 a.u. and a range of radii from 8.4 a.u. to 21 a.u. In addition, these features are not evenly distributed in the space and take up only 58.19% of the space between the spherical angles of 0.24π < θ < 1.8π and 0.21π < φ < 0.95π. This may suggest that the features that are important for capturing human similarity judgments of concepts are a small subset of all features present in object images, or that the experimental methods that generated the 49 features in the THINGS database do not fully capture the features that are important to object image similarity.

**Fig. 4:**
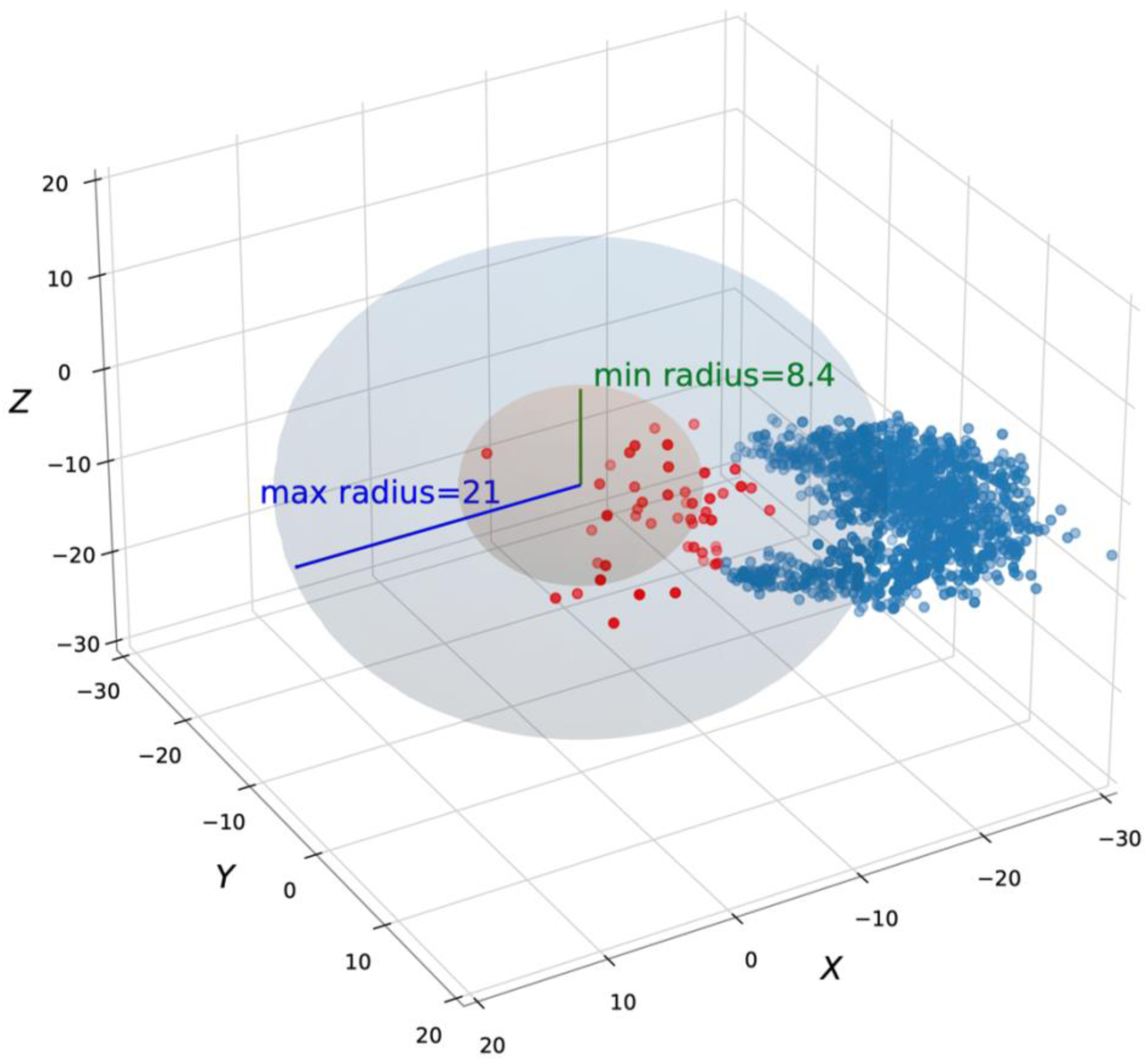
The minimum (8.4 a.u.) and maximum radii (21 a.u.) of the 49 latent features. Red points represent the 49 latent features in the Hyperbolic space with a maximum radius of 21 a.u.. Blue points represent the 1,854 object concepts.

Since the 49 features are not evenly distributed in the space, the 1,854 object concepts which are linear combinations of the features are also not evenly distributed in the space (Fig. 4). Note that coordinates of individual object concepts often fall outside of the maximum Hyperbolic radius of the feature space because the 49 concept features explicitly specify non-negative feature scores for each concept [39]. The concepts form three protrusions along a curved, sphere-like surface which is concave towards the origin.

We also observed that concepts appear to be more clustered by their category membership in the 3D Hyperbolic space compared to the first three dimensions of the 16D Euclidean space (Fig. 5). To verify this observation, we computed a within-between distance ratio to describe the clustering of each category; a ratio greater than 1 means that concepts in this category are more spread out and closer to other categories than to each other. We started with the Euclidean within-between distance ratio, which is the ratio of the average Euclidean distance between concepts (e.g., cat and snake) in the same broader category (e.g., animal) to the average Euclidean distance between concepts (e.g., cat and taco) from different categories (e.g., animal and food; see Methods). Comparing the Euclidean within-between distance ratios from the 16D Euclidean space (M=0.71, SD=0.15) and from the 3D Hyperbolic space (M=0.64, SD=0.14), we indeed find that categories tend to be better separated from each other in the 3D Hyperbolic space compared to the Euclidean space, where smaller ratios indicate better separation (t_paired_= −3.6, *p* < .002).

**Fig. 5:**
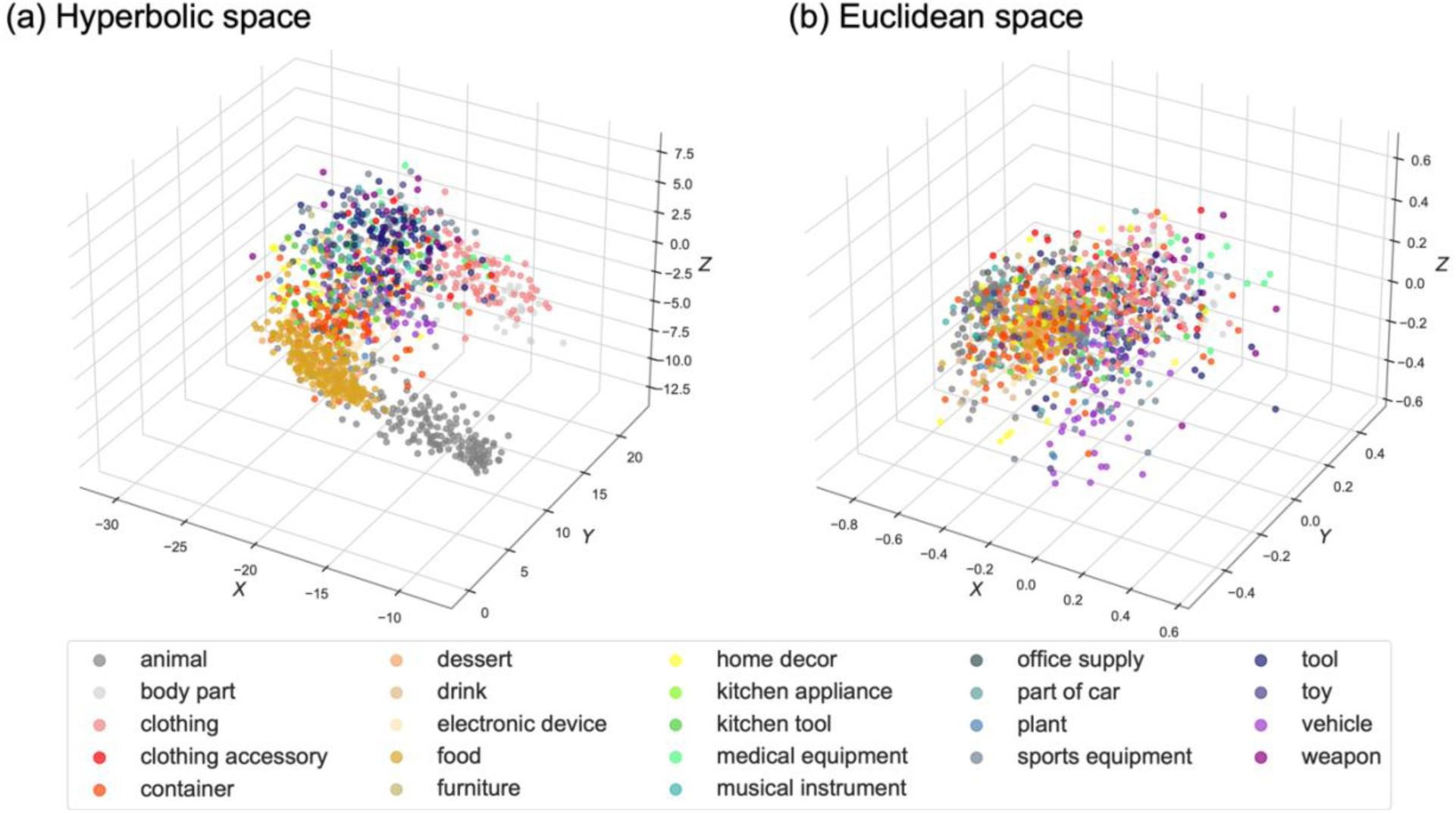
Concepts in the same categories clustered together more in 3D Hyperbolic space than in 16D Euclidean space. **(a)** Categories in the 3D Hyperbolic space. **(b)** Categories in the first 3 dimensions of the 16D Euclidean space.

However, we found that all categories have Euclidean distance ratios smaller than 1 (i.e., all categories are generally more clustered than spread out) in both spaces. To more accurately capture the arrangement of categories in the Hyperbolic space, we computed the Hyperbolic within-between distance ratio (M=0.97, SD=0.07). We found 10 categories with Hyperbolic distance ratios greater than 1, meaning that these concepts have shorter Hyperbolic distance to other categories than to their own categories (Supplementary Fig. 1). These categories, such as *clothing accessory*, *furniture*, and *sports equipment*, are more specialized than those with low ratios, such as *animal*, *food*, and *tool*. This suggests that the Hyperbolic geometry arranges concepts differently and better differentiates specialized categories from general ones than the Euclidean geometry does.

Indeed, the level of clustering of each category is closely tied to its spatial location in Hyperbolic space, which can be measured by the radius from the origin of the space. We found a moderate positive relationship between average concept radius in each category and the Hyperbolic within-between distance ratio in Hyperbolic space (ρ = 0.4, *p* < .001, Supplementary Fig. 2), indicating those general categories that cluster tightly are situated closer to the origin. On the contrary, the within-between Euclidean distance ratio is weakly and negatively correlated with radius in Euclidean space (ρ = −0.08, *p* < .001, Supplementary Fig. 3), meaning that categories further away from the origin are slightly more clustered.

This spatial organization in Hyperbolic space can be observed from example object categories. Concepts from more general categories (e.g., *animal* with an average radius = 19.47 and *food* with an average radius = 19.71) have lower average radii, and these categories have within-between distance ratios smaller than 1, indicating shorter distances within these categories than from other categories. In contrast, concepts in more specialized categories (e.g., *home decor* with an average radius = 26.30 and *clothing accessory* with average radius = 26.03) have longer radii, and these categories have within-between ratios greater than 1. This indicates that, for example, the Hyperbolic distances between different pieces of *home decor* concepts (e.g., couch and mirror) are greater than the distances between these concepts and concepts of other categories (e.g., concepts in *home decor* vs. concepts in *food*). In other words, more general categories (e.g., *food* and *animal*) are more tightly clustered together in 3D Hyperbolic space compared to more specialized categories (e.g., *home decor*), which are more spread out.

These effects were not observed using Euclidean geometries. Compared to concepts in more general categories (e.g., *food* with an average radius=1.02 and *animal* with an average radius=1.06), those in more specialized categories (e.g., *clothing accessory* with an average radius=1.03 and *home decor* with an average radius=1.00) are similarly close to the origin of the Euclidean space, while all categories are more clustered than spread out (i.e., Euclidean within-between distance ratio smaller than 1). This means that 3D Hyperbolic spaces set up hierarchies of semantically related categories where more general categories are closer to the root of the tree and are more tightly clustered together, while more specialized categories are further from the root and are more spread out.

To clearly illustrate these characteristics in this Hyperbolic space, we highlight some key example points within the space. In Fig. 6, concepts in the *body part*, *clothing*, and *clothing accessory* categories (“body part grouping”) are adjacent to each other. The *body part* category, which may be considered a more general category, is closest to the origin (avg r = 22.88) and most clustered (Hyperbolic within-between ratio = 0.92). Within the *body part grouping*, concepts like *knee* (r = 17.39) and *torso* (r = 18.58) are closer to the origin than concepts like *eye* (r = 27.67) and *tooth* (r = 29.30). Concepts related to c*lothing* (avg r = 24.11) are slightly farther from the origin than *body part* and more spread out (Hyperbolic within-between ratio = 0.96), although their radii are not significantly different from the latter (t(141) = 1.97, *p* = .051). Finally, *clothing accessory* is farthest (avg r = 26.03) compared to the two other categories (t_body_(47) = 3.16, *p* = .003; t_clothing_(122) = 2.18, *p* = .031) and is the most spread out (Hyperbolic within-between ratio = 1.04), demonstrating how this category is more specialized. Thus these three related categories are organized along a rough semantic hierarchy that is naturally captured in Hyperbolic space, where *body part* is closer to the root of the network, *clothing* is at an intermediate distance, and *clothing accessory* is farthest away from the root. Such a fine-grained organization of the semantics of these categories cannot be found in the 16D Euclidean space (Fig. 5b).

**Fig. 6:**
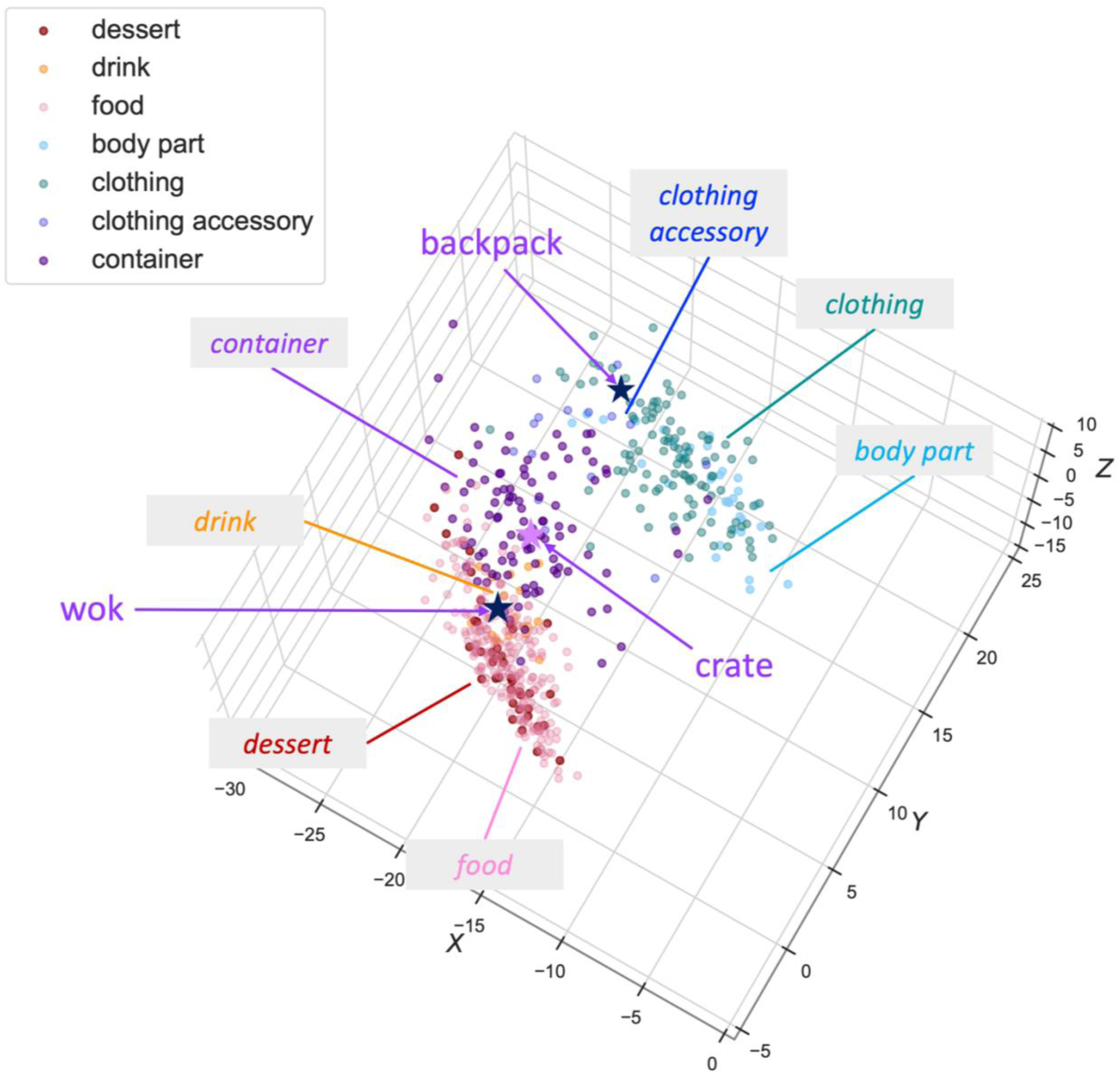
Hierarchical organizations are formed by semantic-related categories in Hyperbolic space. *Body part*, *clothing*, and *clothing accessory* form a tree-like *body part grouping* (upper right). *Food*, *dessert*, and *drink* form a hierarchical *food grouping* (lower left). These two groupings are connected by the *container* category (purple), in which concepts like “backpack” and “wok” (dark purple stars) are situated close to semantically-related categories, whereas “crate” (light purple star) sits far away from other categories.

A similar hierarchical organization can be found within the *food*, *drink*, and *dessert* categories (“food grouping”), in which the *dessert* category (avg r = 19.36) is closely mixed with the *food* category (avg r = 19.71) and both are tightly clustered (Hyperbolic within-between ratio = 0.84 for both), while the *drink* category (avg r = 21.32) is furthest from the root of the network (t_dessert_(39,18) = −2.69, *p* = .009; t_food_(239,18) = −3.64, *p* = .001) and is most spread out (Hyperbolic within-between ratio = 0.89). These two semantic hierarchies, the *body part grouping* and the *food grouping*, are connected by the *container* category. This broad category group of containers includes concepts that bear characteristics of multiple categories. Containers that are specific to food or clothing sit on the boundaries of the *body part grouping* and *food grouping*. For instance, “backpack” is a concept in the category of *clothing accessory*, while it functions as a container and thus is located close to the category of *container*. On the other hand, “wok” is a container for food, which can explain why it is placed at the boundary of categories of *container* and *food* (Fig. 6). In contrast, containers that are non-specific are in the central mass of the *container* category and away from the *body part grouping* and *food grouping* boundaries (e.g., “crate”). Unlike in the 16D Euclidean space where spatial locations are hard to interpret, these meaningful concept geometries emerge naturally from the Hyperbolic embedding of the THINGS feature similarity matrix.

### Concepts closer to the origin are more memorable

We now return to our central question of what the underlying geometry of these feature spaces can reveal about object image memorability. Memorability scores (corrected recognition (CR) scores, obtained by subtracting the false alarm rate from the hit rate for each image) were collected in prior work using the 26,107 object images in THINGS, in which, on average, 40 participants per image (N = 13,946) detected repeated object images in a continuous visual recognition task [10]. These scores are averaged across all exemplar images for each concept to obtain a concept-level memorability score (Mean = 0.79, min = 0.65, max = 0.91, SD = 0.04).

We first focus on whether the radius of each concept can predict its memorability. Although various sets of category-specific semantic or visual features have been examined to predict memory in previous studies [10,12,14,17,40], we aim to uncover mechanisms of memorability that can be generalized across all feature spaces. The radius of each concept, reflecting its relative location (i.e., distance) from the origin of a feature space, is an invariant measure across different features and thus can reveal an underlying principle of what makes something memorable.

We discover that radius and memorability are significantly negatively correlated in both Euclidean and Hyperbolic embedding spaces (Spearman’s Rank Order Correlation, 3D Hyperbolic: ρ = −0.21, *p* < .001, R^2^ = .037; 16D Euclidean: ρ = −0.10, *p* < .001, R^2^ = .009), indicating that concepts closer to the origin, or the center of the representational spaces, are more likely to be remembered across people. Interestingly, the radii in these two spaces are negatively correlated (ρ = −0.14, *p* < .001), suggesting that while these two spaces capture similar aspects of object concepts, they arrange them differently (Supplementary Table 2). This may explain our observation that the semantic hierarchies that emerge naturally in 3D Hyperbolic space are not preserved in 16D Euclidean space. Importantly, we note that radius explains approximately four times more variance in memorability scores in the 3D Hyperbolic space compared to the 16D Euclidean space, demonstrating that the spatial organization of object concepts in the Hyperbolic space tells us more about object memorability.

### Typicality predicts object memorability

In addition to memorability, concept radius in both Hyperbolic and Euclidean spaces may also capture typicality, which is related to memorability in prior works (e.g., [10]). Therefore, we next tested measures of typicality that are more similar to those used in previous studies. We employed two measures of concept typicality that describe the spatial relationship of concepts in categories (Fig. 7): within-category attraction and between-category repulsion. We defined *within-category attraction* using the within-category distance between concepts. That is, the average distance between a concept and other concepts in the same category while accounting for the variation of Hyperbolic distance between different categories (see Methods). This measure captures whether a concept is closer to or farther from other concepts in the same category relative to the average distance between concepts within that category. Concepts with higher within-category attraction have shorter within-category distance, are closer to other concepts in the same category, and are conceptualized as more prototypical to their categories. Within-category distance was strongly correlated with radius in the 3D Hyperbolic space (ρ = .81, *p* < .001, R^2^ = .653), suggesting that those farther away from the origin have longer distances from their own categories and thus show weaker within-category attraction. The variance inflation factor between the attraction measure and radius in the Hyperbolic space was small (VIF = 1.022) indicating that a linear model would be able to statistically identify independent effects of radius and within-category attraction. However, within-category distance was weakly and negatively correlated with radius in the 16D Euclidean space (ρ = −.06, *p* =.03, R^2^ = .015).

**Fig. 7:**
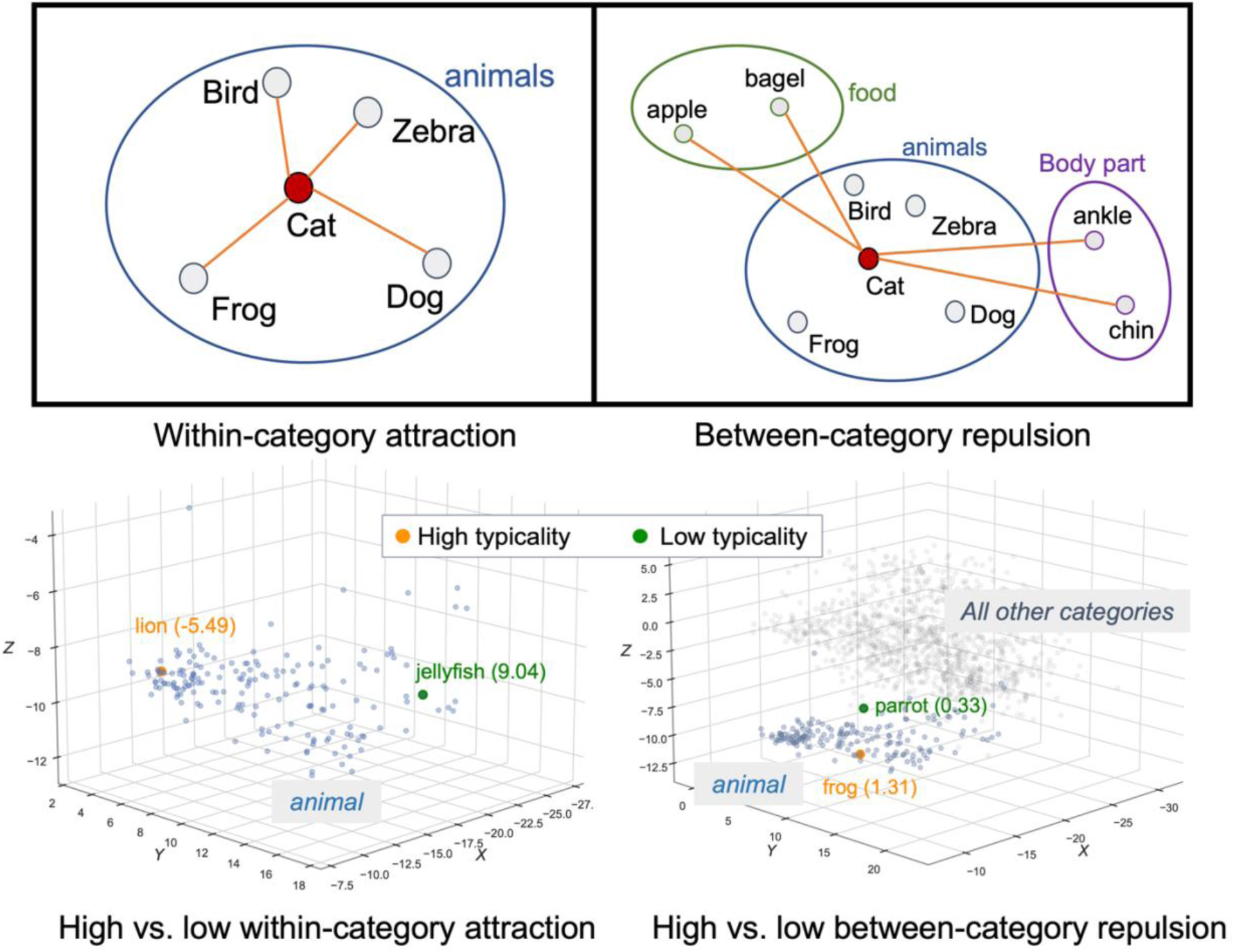
Two measures of typicality (top row) and examples with high vs. low scores in each measure (bottom row). Within-category attraction captures the proximity of a concept to other concepts in its category; for example, lion (within-category distance = −5.49) is much closer to other concepts in the animal category than jellyfish (within-category distance = 9.04) is, and thus the former has a stronger within-category attraction. Between-category repulsion highlights how far a concept is from concepts in other categories; for instance, parrot (between-category distance = 0.33) is closer to concepts outside of the animal category than frog (between-category distance = 1.31) is, and thus the latter has a stronger between-category repulsion.

Our second typicality measure, *between-category repulsion*, is designed to capture our observation that in the Hyperbolic embedding, semantically related categories are connected by concepts sharing characteristics of both categories (e.g., in Fig. 6, backpack and wok connect categories of *clothing accessory*, *container*, and *food*). As a result, there may be some object concepts that are similar to multiple broader categories, while other concepts (e.g., crate) may only relate to their own category (e.g., *container*). While within-category attraction captures how similar concepts are to their own category, between-category repulsion captures how distinct concepts are from other categories. Specifically, we defined between-category repulsion by starting with the average distance between a concept and concepts in other categories and then regressing out radius across concepts (see Methods). These relative metrics rule out the influence of expanded distances at larger radii in the Hyperbolic space. Confirming that regressing out radius results in a relative measure, we found no correlation between radii and between-category repulsion in both Hyperbolic and Euclidean spaces (3D Hyperbolic: ρ = 0.039, *p* = .15, R^2^ = .007; 16D Euclidean: ρ = −.02, *p* = .6, R^2^ = .195).

Including these two typicality measures and the radius of each concept in an ordinary least squares regressions to predict memorability, we find a similar pattern in the Hyperbolic space associating higher memorability with: (1) lower radii (β = −0.03, *p* = .007), (2) higher within-category attraction (i.e., shorter within-category distance) (β = −0.04, *p* = .027), and (3) higher between-category repulsion (β = 0.59, *p* < .001). This indicates that memorable concepts tend to be more prototypical, both globally (closer to the origin) and locally (more typical within a category). However, there is also some evidence for distinctiveness, as higher between-category repulsion was a significant predictor of higher memorability. This means that concepts that are more distinct from concepts in other categories are better remembered. For example, a *lion* is a typical animal and is close to the origin (radius = 14.02, within-category distance = −5.49, between-category repulsion = 0.71, memorability = .83). Furthermore, concepts that are more different from other categories also tend to be more memorable (e.g., a *jellyfish* is an animal that is far from other categories (radius = 27.9, within-category distance = 9.05, between-category repulsion = 1.52, memorability = .81).

Moreover, we find that radius, within-category attraction, and between-category repulsion together explain more variance in memorability when calculated in the 3D Hyperbolic space (R^2^ = .153) compared to the 16D Euclidean space (R^2^ = .061) (Fig. 8). In a previous study using the same THINGS stimuli in a Euclidean framework [10], 38.5% of the variance in memorability was captured by all 49 semantic and visual features, which was calculated to be approximately 61.7% of the noise ceiling (i.e., the maximum proportion of variance that could be achieved by a theoretically perfect model). Here, using only three predictors, we show that global geometry (radius) and local geometry (between and within typicality) of object image feature space capture 15.3% of variance in object image memorability, which translates into 24.5% of the noise ceiling. Importantly, these three predictors can be calculated for any stimulus space, in contrast with visual or semantic features that are specific to a given stimulus category (e.g., the brightness level of object images). Specifically, based on the co-occurrence of features in any stimulus set, we can translate the feature space into a Hyperbolic geometric space for the stimuli and calculate these three predictors. Additionally, conducting analyses in a Euclidean framework alone results in the conclusion that the geometry of object image space explains less than 10% of the variance in object image memorability. This suggests the importance of feature space geometry in memory research, and that the Hyperbolic framework can provide more insights in understanding the drivers of stimulus memorability.

**Fig. 8:**
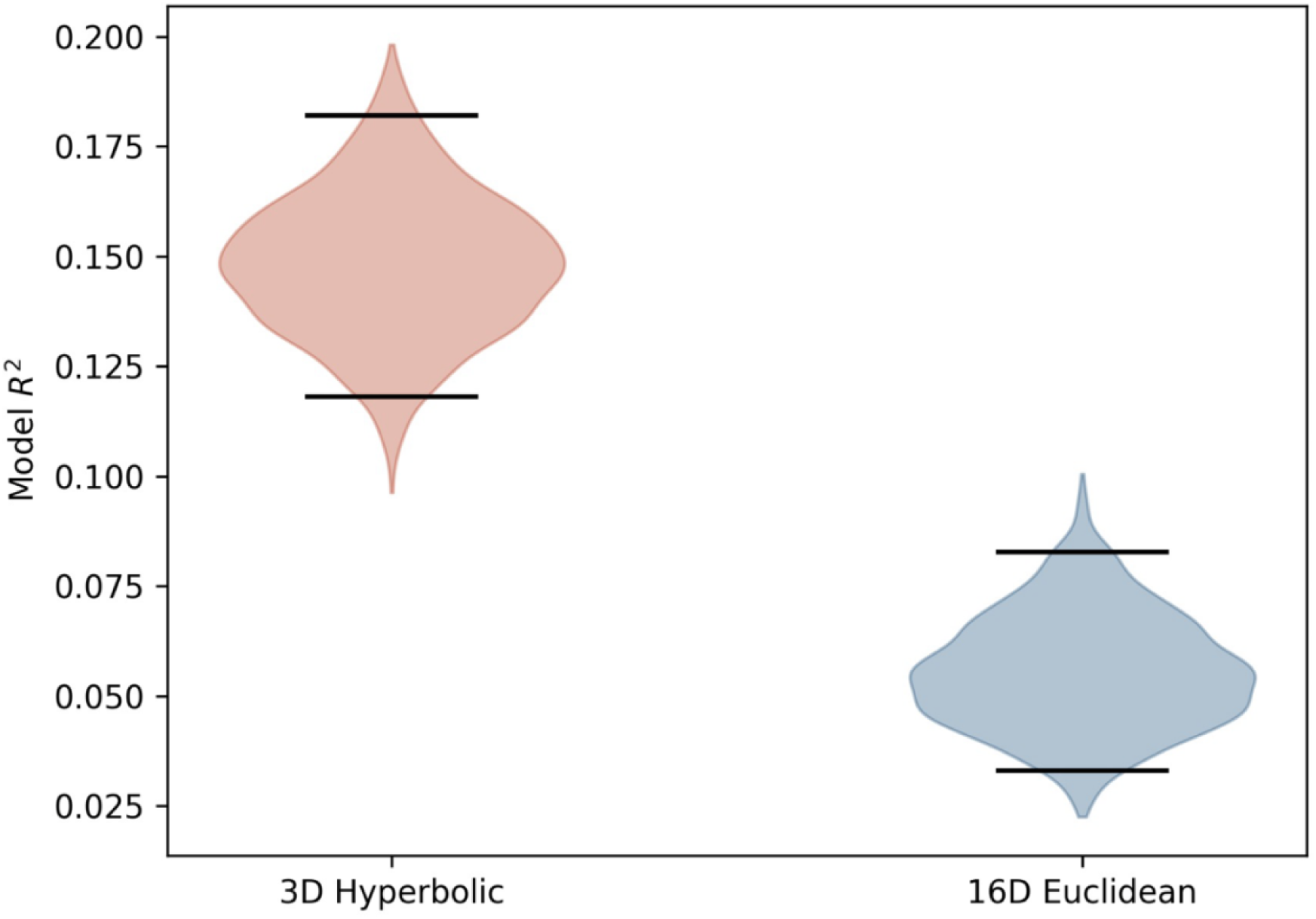
Predictions of memorability using radius, within-category attraction, and between-category repulsion. Predictions of memorability were generated from leave-one-out cross-validation, omitting one concept at a time. Bootstrapped R^2^ values were subsequently computed by resampling these memorability predictions with replacement. A 95% bootstrap confidence interval is indicated by the black bars. The confidence intervals for the two spaces show little overlap, suggesting that memorability is better explained with these three predictors in Hyperbolic space than in Euclidean space.

## Discussion

In this study, we examined how memorability arises in concept space by comparing Euclidean and Hyperbolic geometries. Compared to the Euclidean geometric framework that has been conventionally adopted by most prior research, a curved and hierarchical Hyperbolic space model captures intuitions of object images and their semantic relationships with fewer dimensions and a better fit to the empirical data. Furthermore, Hyperbolic space explained concepts’ memorability better in relation to their spatial locations than the conventional Euclidean geometry. Within the Hyperbolic space, we found that memorable concepts tend to be closer to the center of the entire space, and also more central/prototypical of their own category. As expected, typicality also shows a negative correlation with the distance of concepts from the center, supporting our hypothesis that prototypical concepts are closer to the center of the space. In addition, we found that memorability was related to two measures of prototypicality (within-category attraction and between-category repulsion) that captured the local geometry of object concepts. Finally, using a combination of these global (distance to center) and local (typicality) geometric measures within a Hyperbolic geometry, we were able to explain a substantial amount of the variance in memorability, and more than double the variance explained by a Euclidean space. These findings demonstrate the importance of considering alternate representational spaces, and reveal new generalizable principles about memorability that can cut across stimulus domains.

We present a novel finding about human memory by showing that Hyperbolic geometry is a more suitable model for object memory space, and that object concept representations are organized in hierarchical structures. While prior research has been navigating object memory space in Euclidean frameworks [10,12,13,17,18,19,20], our study demonstrates that the flat Euclidean geometry does not fully model the organization of object concept representations. Instead, the Hyperbolic space model better fits the empirical dissimilarity among the 49 object image features from THINGS with fewer dimensions (3 dimensions) than a Euclidean space model (16 dimensions). Moreover, the inherently hierarchical structure of Hyperbolic geometry captures a greater extent of clustering and distinction among concepts. Specifically, concepts in more general categories cluster closer to the roots of the hierarchically organized network (i.e., the Hyperbolic space), while simultaneously distinguishing concepts in more specialized categories with increased distances. In addition, the Hyperbolic space demonstrates the fine-grained semantic relationships among categories and concepts through hierarchical organizations, which is not observed in the Euclidean space. The systematic patterns of object concepts and categories we found only in the Hyperbolic space challenge the convention of how object concept representations are organized and could revolutionize the geometric framework in which object memory space operates.

By reevaluating the spatial arrangement of object concepts in memory space, we have offered novel perspectives on object memorability. While prior studies proposed that memorability may relate closely to the characteristics of stimuli, here we illustrated that the geometric organization of stimuli within semantic space can drive some of the memorability distinction among items. Our results suggest that memorability can be better explained in a curved Hyperbolic space than in the traditional flat Euclidean space. Within this geometric framework, our study also adds evidence and support to the literature on the originally elusive relationship between typicality and memorability. Specifically, our data in both Hyperbolic and Euclidean representation spaces agree with prior memory research that memorable stimuli are placed closer to the center of the space [8,14], such that stimuli with a shorter distance from the origin have a higher memorability score. We then showed a positive relationship between within-category attraction and memorability, suggesting that memorable items share a greater similarity in their feature representations (i.e., they are more prototypical category members), which is also aligned with previous work (e.g., [8,14]). Moreover, as many previous studies measured stimulus typicality only by how similar they are to their own categories (the “within-category attraction” in our study) and found mixed results on the relationship between memorability and typicality [10,17], our investigation of the between-category repulsion metric provided a different aspect of typicality. We found that memorability is largely influenced by the distance between a concept and all other categories to which it does not belong. In other words, it seems that memorable items may be those that most clearly belong to a single category (i.e., a measure of distinctiveness), and serve as prototypical examples of that category. In this case, the most memorable items are both prototypical (within category) and distinctive (between category).

While our findings provide new insights into visual memory and the object concept space, there are several potential directions for future study. First, each dimension in this new representation space no longer maps onto specific features but rather organizational aspects of those features (e.g., radius). This is different from many prior works in which features serve as the dimensions of the representational space of a set of stimuli, and the feature embeddings of each stimulus determine its coordinates in the space [12,14,40]. Instead, by moving into a lower dimensional space via the Hyperbolic multidimensional scaling (MDS) process, the three dimensions now no longer represent any direct properties of the features and concepts, but instead capture broader organizational principles such as a concept’s distance from the origin. As a result, our findings can be generalized to other datasets beyond object images, even when items in those datasets are defined by embeddings of entirely different features. For example, we hypothesize that this relationship between a shorter radius and memorability could translate to other cognitive domains and stimulus categories, such as words [8] or other sensory modalities (e.g., smells). Such generalizations can be achieved through our ALBATROSS method by applying the process to any set of features and stimuli, through which it can introduce the hierarchical structure and generalize our findings to other cognitive spaces.

Second, the concept spaces we inspected in this study, Euclidean and Hyperbolic, are constructed based on the inferences of human-judged similarity among object concepts. Therefore, they may not capture the entire spectrum of factors that drive memorability and may not unveil the full structure of concepts in the human brain, if features that humans are not consciously aware of also determine this structure. Furthermore, since the features that construct the representational spaces were derived from one exemplar image for each concept, they may not describe all properties of the 1,854 concepts. These aspects of the original dataset may explain why the features are arranged in only one section of the spaces (Fig. 4), and thus we may not be able to explain all of the variance in memorability. Even so, we demonstrated that even these incomplete object concept representations are organized in a hierarchical structure rather than in a flat Euclidean geometry, and that Hyperbolic geometry efficiently captures variance in memorability with fewer dimensions.

While human behavioral data has revealed an underlying hierarchical structure of concepts, we are curious if the same Hyperbolic structure of memory exists in the human brain, which is an exciting avenue for future potential work. Therefore, future studies can utilize neuroimaging data and the same set of object concepts in the THINGS database to model the representation space in the brain and examine how memorability relates to a hierarchical arrangement. If the relationship is consistent with the behavioral data, this finding will be strong evidence for a non-Euclidean memory space geometry from a neuroscientific perspective.

Finally, our dataset on concrete objects theoretically covers all nameable and imageable concrete object concepts. Future work could examine other cognitive spaces, such as the spaces of abstract concepts, actions, or concept spaces built within other cultures or languages. By collecting and placing these concepts in a Hyperbolic space, we can investigate if memorability and typicality show patterns parallel to what we found in the space of concrete object concepts. Moreover, future work could attempt to generate novel items within this Hyperbolic space and examine whether they are remembered differently based on where they fall in the space. Linking to our observation that the 1,854 concrete object concepts formed a skewed distribution in the 3D Hyperbolic space, we speculate that these other types of concepts may take up part of the empty space that the current 49 features do not capture.

Overall, our study reveals the underlying hierarchical structure of concrete object similarity and semantics by showing that a nonlinear Hyperbolic geometry better models the semantic space and subsequent human memory of object concepts than the traditional Euclidean geometry. In this Hyperbolic space, we demonstrate that memorable and prototypical object concepts tend to be situated closer to the center of the space. Finally, our results suggest a positive correlation between concept memorability and typicality. Radius and within-category attraction (both proxies for object typicality/prototypicality) show a positive relationship with memorability, while between-category repulsion (a proxy for distinctiveness) also shows a positive relationship with memorability. Our findings redefine the landscape of visual memory research by presenting an alternative non-Euclidean framework for concept memory space that elucidates object memorability and challenges the conventional assumption of Euclidean geometry. Furthermore, through a topological lens, our study offers a comprehensive understanding of the relationship between memorability and typicality – showing that objects that take the perfect balance of being canonical yet distinctive are the objects that are remembered best.

## Methods

### Data

Our data consists of concrete object concepts and their corresponding memorability scores. We included all 1,854 object concepts (e.g., apple, backpack, cat) from the THINGS database [38], which were sampled as the picturable and namable American English nouns in the WordNet lexical database and theoretically covered most concrete object nouns in English. Each concept has at least 12 naturalistic images as its exemplars, collected through a large-scale web searching process and selected manually from the ImageNet database, forming a corpus of 26,107 object images. Each concept also belongs to one of 27 higher-level categories (e.g., food, clothing, animals) identified by crowdsourced workers and complemented by the WordNet taxonomy. More detail about the database is described in its accompanying paper [38].

Each of the 1,854 concepts is represented by an embedding of 49 features in the THINGS database. Through a triplet odd-one-out crowdsourcing experiment and a data-driven computational model, 49 dimensions were derived from pairwise similarity ratings of all 1,854 concepts and were labeled with a feature name that reflected perceptual properties (e.g., is tall/big) or semantic membership (e.g., is metal/tools) [39]. All feature scores for each concept are non-negative values that specify the extent to which a given feature is represented in this concept.

Memorability scores were collected by Kramer et al. [10] in a continuous recognition paradigm for the 26,107 object images in THINGS. The memorability of each object image was quantified by its corrected recognition (CR) score, which was computed by subtracting the false alarm rate (FAR) from the hit rate (HR) of responses to a stimulus in the visual recognition task. To obtain memorability scores on the concept level, we computed the mean memorability score for each object concept across all its exemplar images.

### Constructing object spaces in different geometries

To compare Hyperbolic space to Euclidean space in explaining memorability of object concepts, we identified the best-fitting models with these two geometries, constructed the representational spaces accordingly, and placed concepts and features into each space. We used ALBATROSS, a stochastic topological analysis technique, to model and construct the Hyperbolic and the Euclidean space that best fitted our data (Stier et al., In prep). Using Pearson correlations between the feature embeddings of concepts from THINGS, we prepared a dissimilarity distance matrix of 49 features, which captured the similarity in occurrences between features. Based on this distance matrix, ALBATROSS modeled and suggested that 16 dimensions in Euclidean geometry can best describe the relationship amongst the 49 features, while only 3 dimensions are required in Hyperbolic geometry. These 3 dimensions reflect radius (distance from the core) and two angle measures.

Next, we placed the 49 features in these two geometric spaces by transforming the dissimilarity distance matrix via dimensionality reduction. In the Euclidean space, we used multidimensional scaling (MDS) to reduce the dimensionality from 49 to 16 (the best-fitting Euclidean model), generating a 49- by 16-dimensional Euclidean feature embedding, which preserves the Euclidean distance between each pair of features. However, the same MDS function cannot be applied to the Hyperbolic space because, unlike Euclidean distance, the distance between two points varies by their distances from the core in the Hyperbolic geometry. Therefore, we used a Hyperbolic version of MDS (Stier A., https://github.com/enlberman/hyperbolicMDS) to scale down dimensionality while preserving the Hyperbolic distance between features. Since the Hyperbolic MDS is a stochastic process, the output from each run was slightly different. To find the optimal locations of features in the Hyperbolic space, we ran the Hyperbolic MDS 500 times and obtained 500 versions of 49- by 3-dimensional Hyperbolic feature embeddings. We then iterated through every version and rotated their orientations using orthogonal Procrustes, which allowed us to align the same feature points across 500 versions to the same location. This mapping process smoothed out the noise among all versions of the 3-dimensional embeddings and generated the final coordinates of each feature in the Hyperbolic space.

Knowing where all features are placed, we located each concept in the spaces. Because each concept is represented by 49 features in the original feature embeddings from THINGS, its location in a space is determined by its unique linear combination of all 49 features. Therefore, we found the location of each concept in the Euclidean space by multiplying the original 49-feature embeddings of concepts with the 49- by 16-dimensional Euclidean feature embeddings, which generated a matrix with 16-dimensional Euclidean coordinates for the 1,854 concepts. Similarly, we obtained 500 versions of concept mappings in the Hyperbolic space by multiplication. Through a Procrustes process, we rotated and averaged these 500 versions of mappings and obtained the final 3-dimensional Hyperbolic coordinates of all 1,854 concepts.

### Category locations and clustering

Each concept belongs to one of the 27 superordinate categories in THINGS. Observing from the category level, we examined different clustering patterns of concepts in the first 3 dimensions of Euclidean space and Hyperbolic space. We computed a within-between distance ratio within each space using the average Euclidean distances between and within each category in both spaces (see equation (1), *n* denotes the number of concepts in one given category, and *m* denotes the number of concepts in each category *c*). We then conducted a paired sample t-test to test whether the clustering of categories in the Hyperbolic space was significantly different from the clustering in the Euclidean space. To more accurately capture the distance between concepts in Hyperbolic space, we also computed the within-between ratio using Hyperbolic distance, and compared it with the within-between Euclidean distance ratio in Euclidean space through another paired sample t-test. Next, to have a more holistic understanding of clustering patterns in both spaces, we examined the Spearman’s Rank Order Correlation between concept radii and within-between Hyperbolic distance ratio in Hyperbolic space, and the correlation between concept radii and within-between Euclidean distance ratio in Euclidean space.

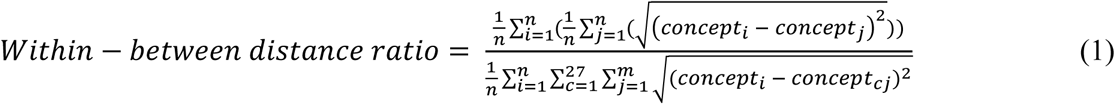

In addition to how close each category clusters, we observed groupings of certain categories that are semantically related and form hierarchies in the Hyperbolic space. To understand whether this hierarchical nature among categories is unique in the Hyperbolic space, we closely investigated these categories and their average radii in both Hyperbolic and Euclidean spaces.

### Typicality measures

Two measures of typicality were computed to capture different aspects of concept typicality, including within-category attraction and between-category repulsion. To obtain the measure of typicality that we dub “within-category attraction”, we computed the within-category distance, which is the mean distance between a concept and all other concepts in the same category, followed by subtracting the average of these mean distances across all concepts in this category. Equation (2) shows the average distance between concept *i* and other concepts *j* in the same category, where *n* denotes the total number of concepts in that category. In equation (3), we ruled out the effect of expanded Hyperbolic distance in different categories by subtracting the mean of average distances of all individual concepts within the category. Specifically, because distances between two points get expanded in Hyperbolic space when their radii increase, concepts in the categories further from the center of the Hyperbolic space may have generally longer distances from their in-category members, leading to an underestimated within-category attraction. Therefore, we accounted for the average within-category distances in each category to identify concepts that have stronger- or weaker-than-average attractions to their in-category members. A concept with a higher within-category attraction has shorter within-category distance and is more similar to most concepts in the same category, whereas a concept far apart from its in-category members has longer within-category distance and lower attraction.

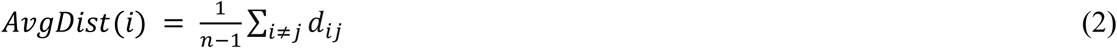

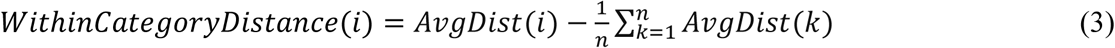

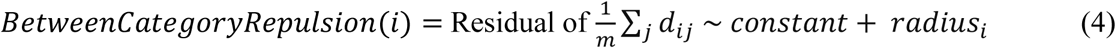

Unlike the within-category attraction measure, the “between-category repulsion” measure captures how far a concept is from other categories in the representational space. To obtain the between-category repulsion of each concept, we started with computing its average distance to all out-category concepts, regressed out its radius, and retrieved the residuals as the final repulsion score (as seen in equation (4), *m* denotes the number of concepts in categories different from concept *i*). Similar to our consideration for within-category attraction, the distances between a concept and out-category concepts can be exaggerated when they have longer radii in the Hyperbolic space, and thus we used residualization to tease out this influence. A concept further from out-category concepts has a higher between-category repulsion score, whereas a concept closer to those outside of its category scores lower in repulsion.

### Analysis of radius, typicality, and memorability

To test our hypothesis that Hyperbolic space can explain concept memorability, typicality, and the relationship between them, we inspected each Spearman’s Rank Order Correlation between these factors across the 1,854 concepts. We first analyzed the relationship between the radii of concepts and their memorability in both Euclidean and Hyperbolic spaces, followed by the relationship between radii and typicality. Instead of analyzing the Cartesian coordinates of concepts in each space, which provide less robust information about concepts, we obtained the radius of each concept by measuring the distance from the origin of the space. Analyses of the concept radius allow us to draw more generalizable conclusions about how memorability changes at different spatial locations, regardless of the concept and feature corpus we used for constructing the representational spaces.

We then ran a series of ordinary least squares (OLS) linear regression models in each space to investigate the relationships among spatial locations of concepts, typicality, and memorability. We started with the first OLS model to predict memorability using concept radius. To examine where prototypical concepts were located in each space, we ran two OLS models using concept radius to predict two typicality measures, within-category attraction and between-category repulsion, respectively. Next, to understand the role of typicality in explaining memorability, we ran two other OLS models testing how each typicality measure predicts memorability. Finally, we took both concept radius and each typicality measure to predict memorability.

## Acknowledgements

This work was funded by the National Eye Institute grant (R01-EY034432) and a Scialog grant (Research Corporation for Science Advancement) to W.A.B.; and National Science Foundation under NSF S&CC-IRG grant #1952050 to M.G.B.

## Supplementary Materials

**Supplementary Table 1.** Hyperbolic space with different dimensionality. Each Hyperbolic space model is compared to the original relationship among 49 features (i.e., empirical data) and the best-fitting 16-dimensional Euclidean model via a chi-square test, with p-values and chi-square statistics reported in the table. A p-value less than .05 indicates that the given Hyperbolic or Euclidean space model is statistically different from the empirical data. The Hyperbolic spaces with 3, 4, or 5 dimensions are not significantly different from the empirical data but start to diverge at 6 or more dimensions, with 3 dimensions being the best-fitting Hyperbolic space model (it has the lowest X^2^).

| Dimension | $p$ (*Hyperbolic*) | $X^2$ (*Hyperbolic*) | $p$ (*Euclidean*) | $X^2$ (*Euclidean*) |
| --- | --- | --- | --- | --- |
| 3 | 1 | 0 | 0.40533943 | 12.513888 |
| 4 | 0.99670414 | 2.81695299 | 0.894859 | 6.39530991 |
| 5 | 0.29409701 | 14.1043083 | 0.77555486 | 8.12162688 |
| 6 | 0.00615747 | 27.681234 | 0.24526226 | 14.9304455 |

**Supplementary Table 2.**
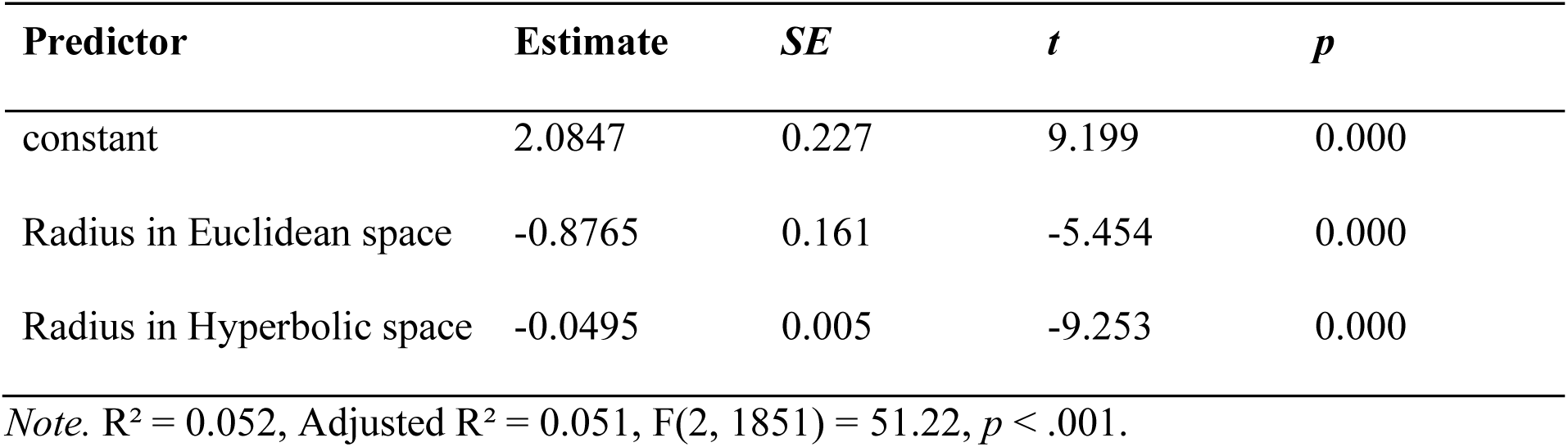
Predicting memorability in an OLS regression model using radii in 16D Euclidean space and 3D Hyperbolic space. Note that the R² of this model is greater than the sum of R² in the two single predictor regressions, where radii in Euclidean (R²=0.009) and Hyperbolic (R²=0.037) spaces predict memorability respectively. With the negative correlation found between the radii from the two spaces, they do not explain any shared variance in memorability.

**Supplementary Fig. 1:**
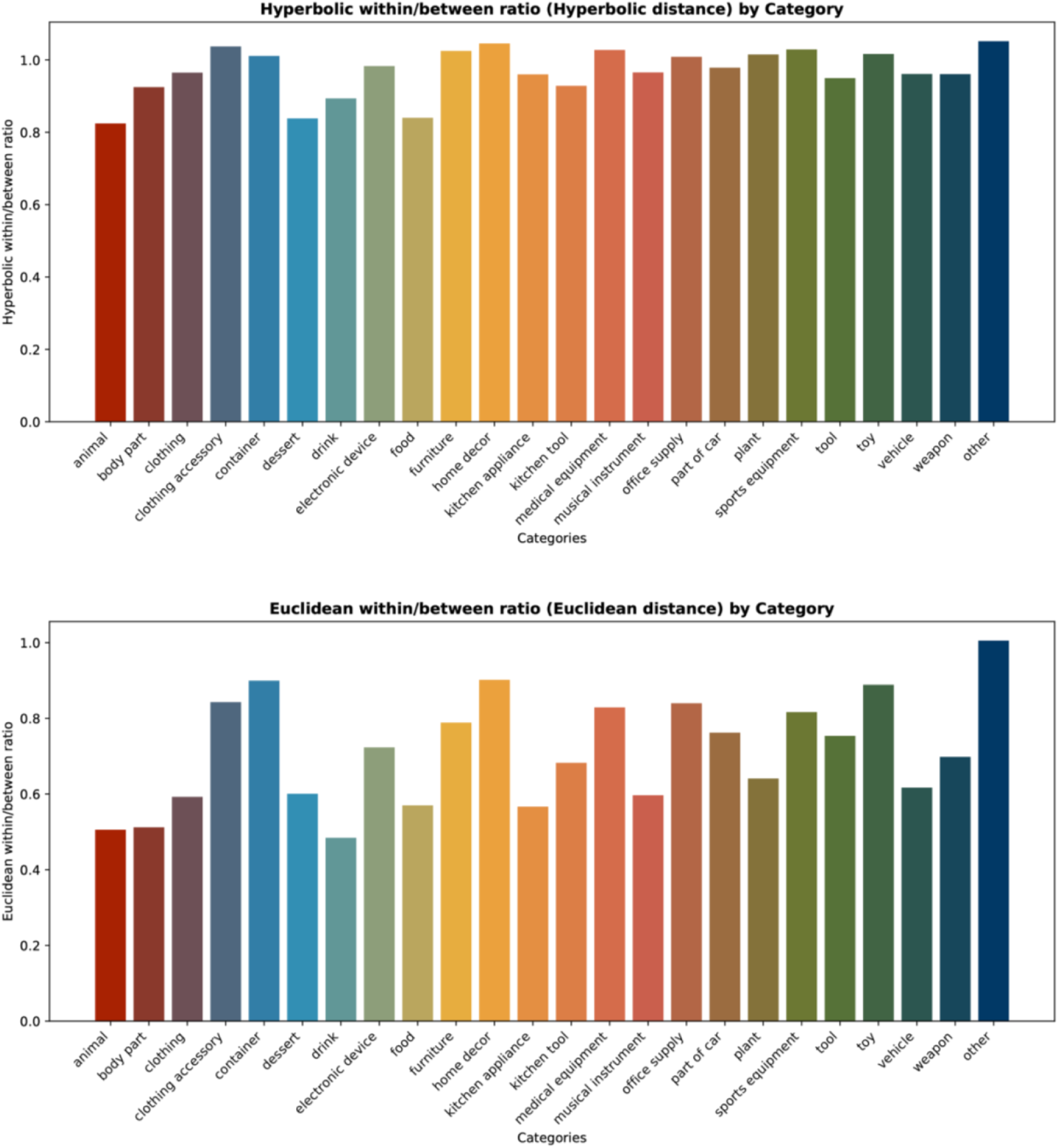
Within-between distance ratio in Hyperbolic space (Hyperbolic distance) and Euclidean space (Euclidean distance). Most categories have a within-between distance ratio closer to 1 in Hyperbolic space compared to in Euclidean space. Moreover, all categories have a within-between ratio smaller than 1 in Euclidean space.

**Supplementary Fig. 2:**
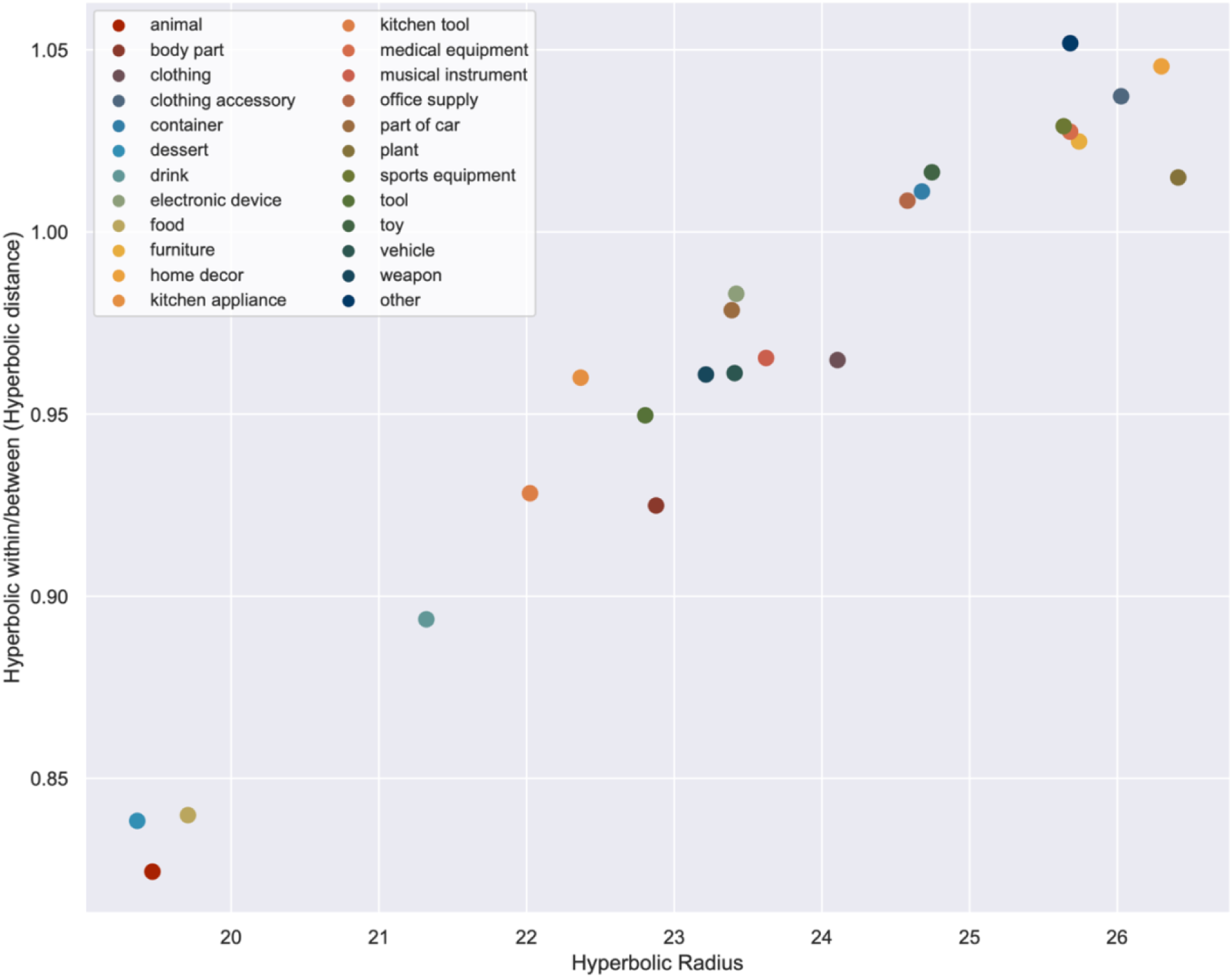
Within-between Hyperbolic distance ratio by different radii in the Hyperbolic space.

**Supplementary Fig. 3:**
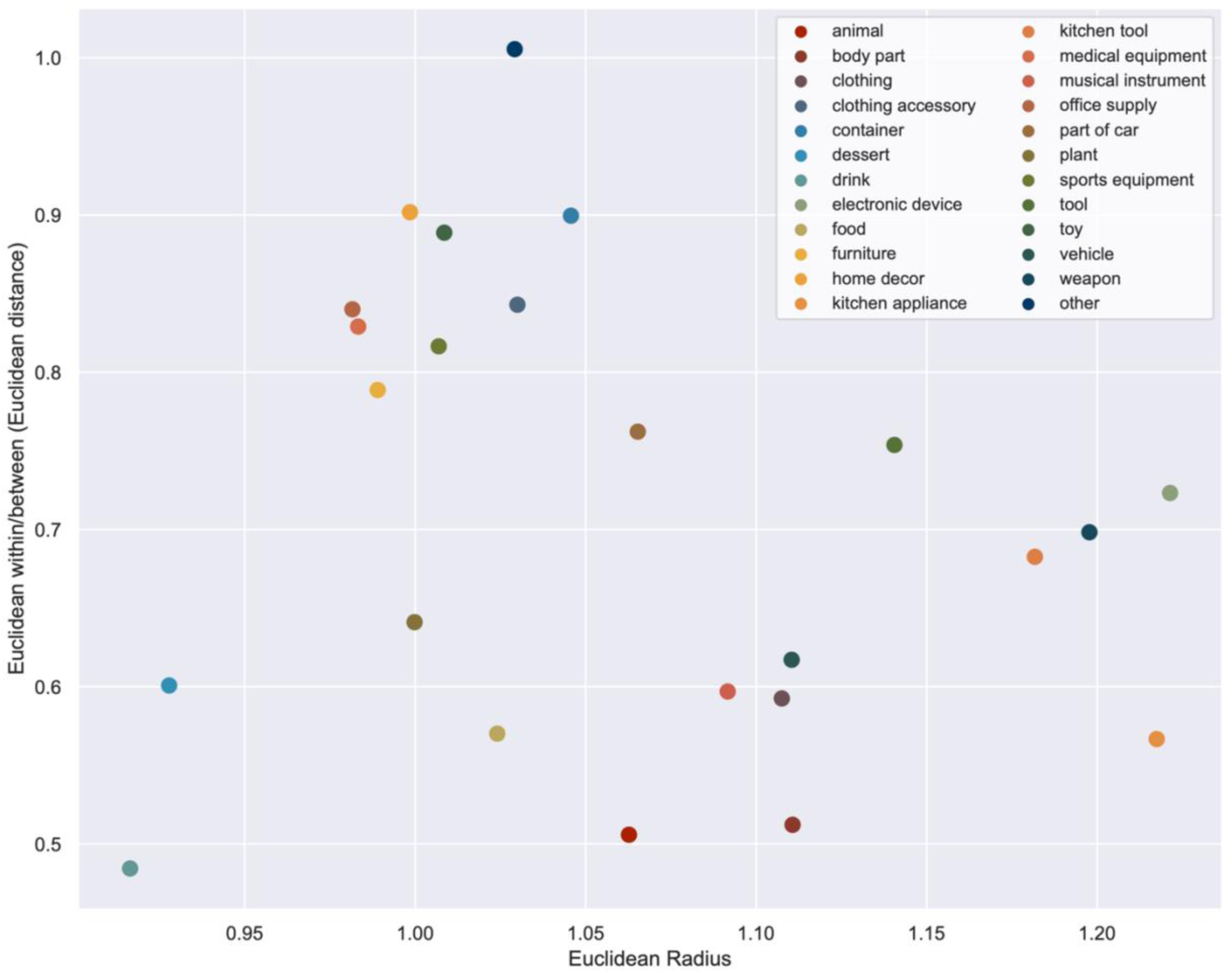
Within-between Euclidean distance ratio by different radii in the Euclidean space.

